# More filtering on SNP calling does not remove evidence of inter-nucleus recombination in dikaryotic arbuscular mycorrhizal fungi

**DOI:** 10.1101/2020.01.15.906412

**Authors:** Eric C.H. Chen, Stephanie Mathieu, Anne Hoffrichter, Jeanne Ropars, Steven Dreissig, Jörg Fuchs, Andreas Brachmann, Nicolas Corradi

## Abstract

We respond to a recent reanalysis of single nucleus sequence data from Chen et al. 2018 *eLife*, which indicated that evidence of inter-nuclear recombination in dikaryotic arbuscular mycorrhizal fungi decreases when heterozygous, duplicated sites being supported by less than 5 reads, are removed from the dataset. Here, we show that applying a more stringent methodology for filtering SNP calls that focuses exclusively on single copy and homozygous regions with at least 5 reads supporting a given SNP, still recovers several hundreds putative inter-nucleus recombination events within the same dataset. We also provide evidence for retaining SNPs supported by less than 5 reads for genotyping individual nuclei using the same dataset.

## Introduction

Genome-based analyses have uncovered a large number of signatures of sexual reproduction in the arbuscular mycorrhizal fungi (AMF), challenging the notion that these organisms are ancient asexuals ^1–7^. In particular, genome analysis showed that model AMF in the genus *Rhizophagus* can be either homokaryotic, carrying thousands of nuclei with a similar genotype, or dikaryotic, whereby nuclei from two parental genotypes are present in the cytoplasm at all times.

Furthermore, the nuclei of dikaryotic AMF isolates each carry one of two divergent regions that resemble the mating-type (MAT) loci of sexual fungi. The MAT-locus is a genomic region that governs sexual identity in fungi ^1^. In dikaryotic sexual fungi, co-existing nuclei are expected to recombine through sex or somatic events ^8,9^. To test the existence of recombination, Chen and colleagues sequenced 86 single nuclei in seven AMF isolates, and showed evidence that rare inter-nucleus recombination events are present in dikaryotic AMF strains ^10^.

A recent comment based on a reanalysis of data from Chen *et al.* 2018 showed that evidence of inter-nuclear recombination in AMF drops significantly once heterozygous, duplicated regions covering SNPs and sites supported by less than 5 reads are removed from the dataset ^11^. Here, we show that the removal of duplicates, heterozygous positions and sites supported by less than 5 reads still retrieves significant inter-nuclear recombination in the dataset provided by Chen *et al.* 2018. Furthermore, by investigating the mapping confidence of single nucleus reads on available genome references, and by validating low coverage genotypes from dikaryotic isolates, we also find little support for the complete exclusion of sites with a low read depth for genotyping individual AMF nuclei.

## Results

### Using stringent filtering methods still recovers hundreds of cases involving inter-nucleus recombination in dikaryotic isolates of Rhizophagus irregularis

Reports of rare inter-nucleus recombination in AMF ^10^ have recently been challenged for having been identified along regions that are, at times, along sites that are heterozygous, duplicated, and with low read depth (< 5 reads supporting a SNP). It was reported that when such sites are removed, the number of inter-nucleus recombination events drops significantly ^11^. We appreciate the utility of a more conservative approach for analyzing single nucleus genome datasets, which are always noisy.

Here, we first aimed to validate the conclusions of Chen *et al.* 2018 using a larger portion of our dataset ^10^. To this end, we tested a stringent method that focuses on sites with a coverage > than 5, and which removes duplicates and heterozygous positions, to identify putative inter-nuclear recombination events across the first 1000 contigs (37% and 50% of the three dikaryotic reference genomes SL1, A4 and A5. Note that the average assembly coverage for single nuclei Illumina read varies from 11% for SL1 to 58% for A5).

We mapped reads from single nuclei against their corresponding dikaryotic reference genomes (e.g. single nuclei reads from SL1 against SL1 reference genome, etc.) and scored SNPs using freebayes with following parameters: -p 1 -m 30 -K -q 20 -C 2.

We consider as evidence of inter-nucleus recombination cases where (1) one or two contiguous SNPs match the haplotype carried by nuclei with the opposite MAT locus (a genomic regions putatively involved in sex determination in AMF); and (2) at least three contiguous SNPs match the haplotype carried by nuclei with the opposite MAT locus.

For scenario #1, which was not analyzed by a recent comment on ^10^, we detected a total of 913 cases (SL1:115; A5:193; A4:605). These mutational events are unlikely to represent sequencing errors or somatic mutations, as they always produce the opposite co-existing genotype (as opposed to random, nucleus-specific substitutions). These sites were recovered along single copy, homozygous sites with at least five reads supporting the given SNP position.

For the scenario #2, where variation along individual single nuclei spans more than three contiguous SNPs, our analysis recovered 195 recombinant blocks (SL1: 36; A5: 30; A4: 129; **Supplemental Table 1, Supplemental Table 2**). This compares to a total of 6 blocks found by others in the smaller portion of the dataset ^11^. These recombinant blocks include up to 17 contiguous SNPs, and between 183 (in the isolate A5) to 635 (in the isolate A4) SNPs in total. Blocks can encompass up to 7 kb of individual nuclear genomes.

This re-analysis of the original dataset published in ^10^ shows that using more stringent filters, i.e. single copy and homozygous sites with at least five reads supporting a SNP, does not remove evidence of inter-nuclear recombination. It also confirms that putative inter-nuclear recombination is rare in AMF, as was originally reported by Chen *et al.* 2018.

### High quality of low coverage sites and recombination in high-coverage sites

Low coverage calls, i.e. positions supported by less than five reads, represent 35% to 54% of available single nucleus data from dikaryotic isolates. High coverage variability is common for single nuclei sequencing data, as it is generated by sequencing DNA subjected to multiple amplification displacement (MDA). Still, the analysis of positions with low-coverage depth, i.e. 2 to 4 reads by Chen *et al.* 2018, was highlighted as a putative source of error in the detection of inter-nucleus recombination, as their removal led to the disappearance of many “recombinant” sites ^11^.

This intriguing correlation begs the question: what causes the link between recombinant sites and low coverage calls? Presumably, low coverage read mapping produces untrustworthy SNP calls.

This assumption conflicts with the nuclear genotypes identified by Chen *et al.* 2018, especially for the two dikaryotic isolates A4 and A5, whose genotypes are said to be mainly produced from positions with low read depth ^11^. Still, despite low read depth, paired-end Illumina reads obtained from their single-nuclei produced a clearly dikaryotic pattern with two nuclear genotypes, each linked with a specific MAT-locus ^10^. This dichotomy should not emerge if the abundant low coverage sites produced false SNP calls in dikaryotic isolates.

To test if the genotyping of individual nuclei is affected by low read depth, we also inspected the mapping quality of single nuclei paired-end reads from 5 homokaryotic isolates (isolates A1, C2, B3 belonging to *Rhizophagus irregularis* species, *Rhizophagus diaphanus* - MUCL-43196, and *R. cerebriforme* - DAOM227022). Importantly, these 5 assemblies carry a size and/or fragmentation that is sometimes larger than those of dikaryotic isolates. This variability in assembly parameters is important because it can negatively affect the quality of SNP calls.

This analysis shows that low coverage calls (i.e. 2 to 4) are all of very high quality in homokaryons, *i.e.* always produces on average the reference genotype > than 99.985% of the time, regardless of size and fragmentation of the assembly (**Figure 1**). Note that the high accuracy of these calls is virtually identical to those made with much larger coverage - *i.e.* from 5 to 100 (99.987%). This shows that low coverage data from single nucleus reads provides an accurate view of nuclear genotypes regardless of assembly statistics, and is thus suitable to genotype individual nuclei.

**Figure 1.**
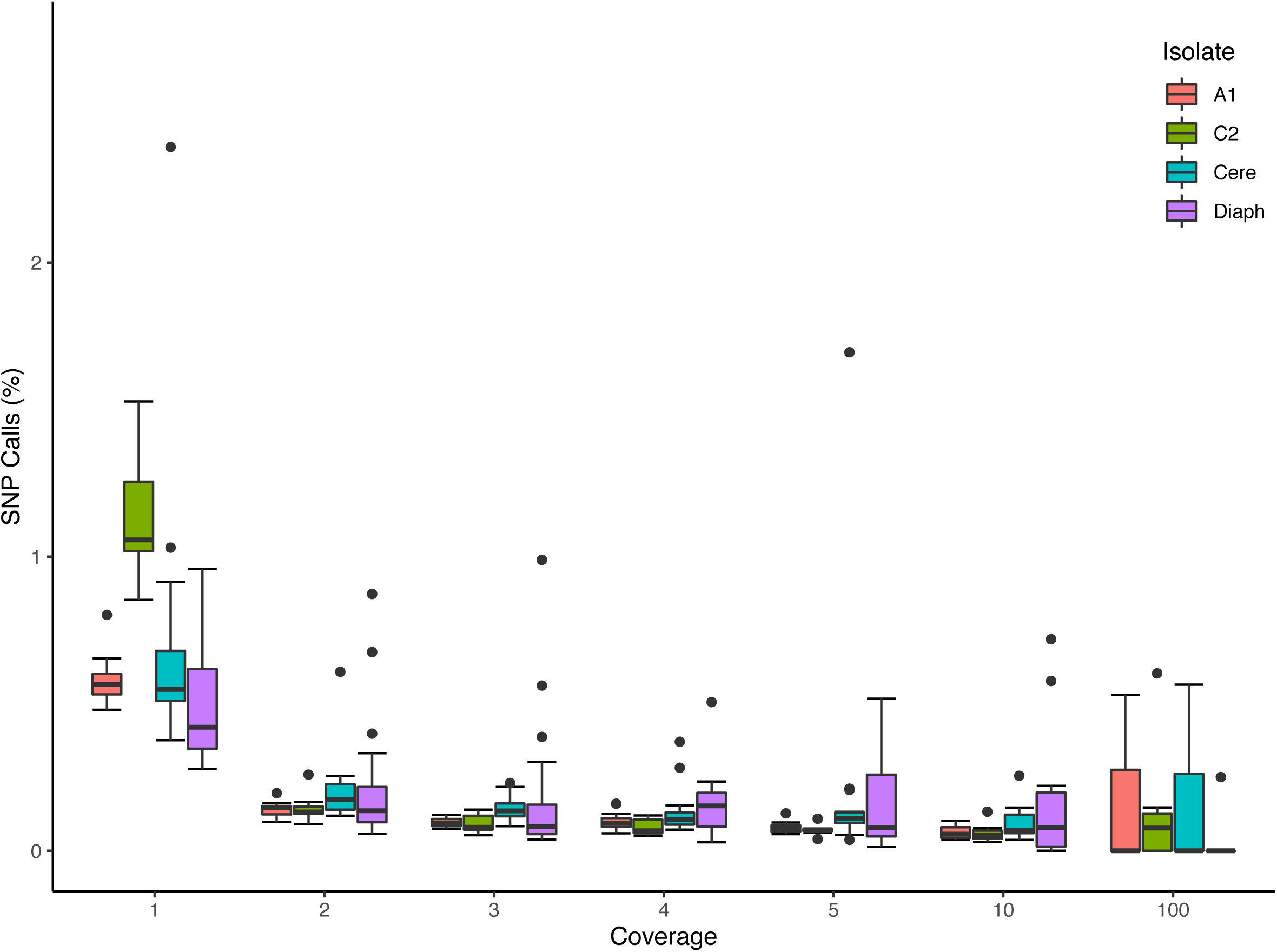
Validation of low coverage depth SNP calls based on single nuclei Illumina reads from four homokaryotic strains. The boxplot represents the percentage of SNPs found to be in disagreement with reference assemblies in two *R. irregularis* isolates (A1 and C2), *R. cerebriforme* (Cere), and *R. diaphanous* (Diaph) organized by the number of reads supporting a given SNP. Boxes represent 25-75% percentile and whiskers represent the largest and smallest value within 1.5 interquartile range above 75^th^ or below 25^th^ percentile. One outlier from *R. diaphanus* is not shown (SN17, coverage 100: 1 mismatch out of 36 positions).

To further support for the retention of low coverage calls in dikaryotic strains, we investigated if low coverage calls – i.e. 2 to 4 reads - could validate a dikaryotic genotype found years earlier using PCR and Sanger sequencing. These genotypes were originally identified from the A5 scaffolds 2, 17, 37 and 641 (see columns representing individual nuclei from A5 in Fig. 3 of Ropars et al. 2016 ^1^).

**Figure 2.**
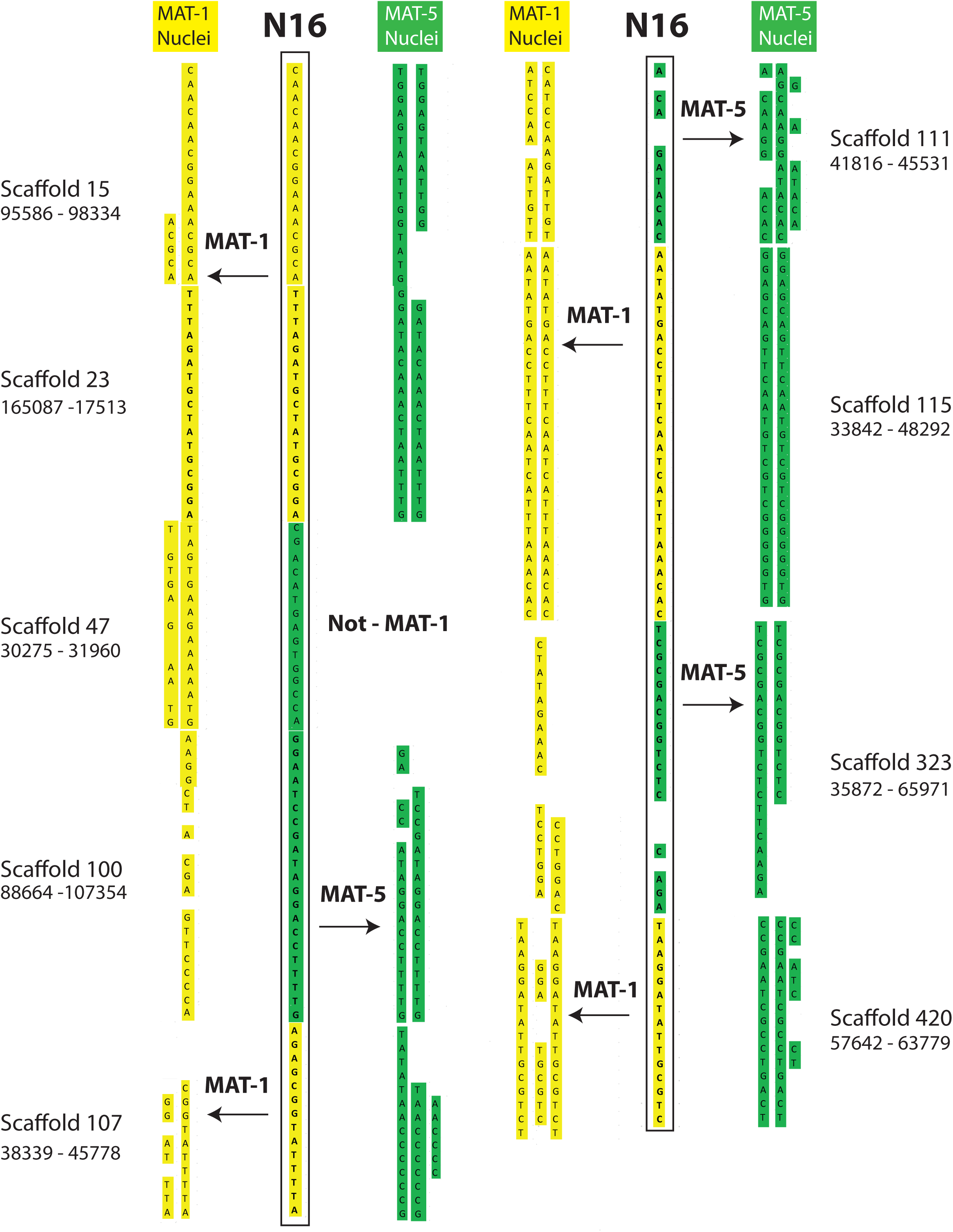
Examples of inter-nucleus recombination in a nucleus of the dikaryotic isolate SL1. The regions are found along homozygous regions present only once in the reference genome of SL1. The nucleus 16 of SL1 carries a genotype that is overwhelmingly similar to nuclei carrying the MAT-1 locus (yellow). In several instances, however, the SN16 is found to switch alleles to carry the other co-existing genotype (green) over several kilobases.

**Figure 3.**
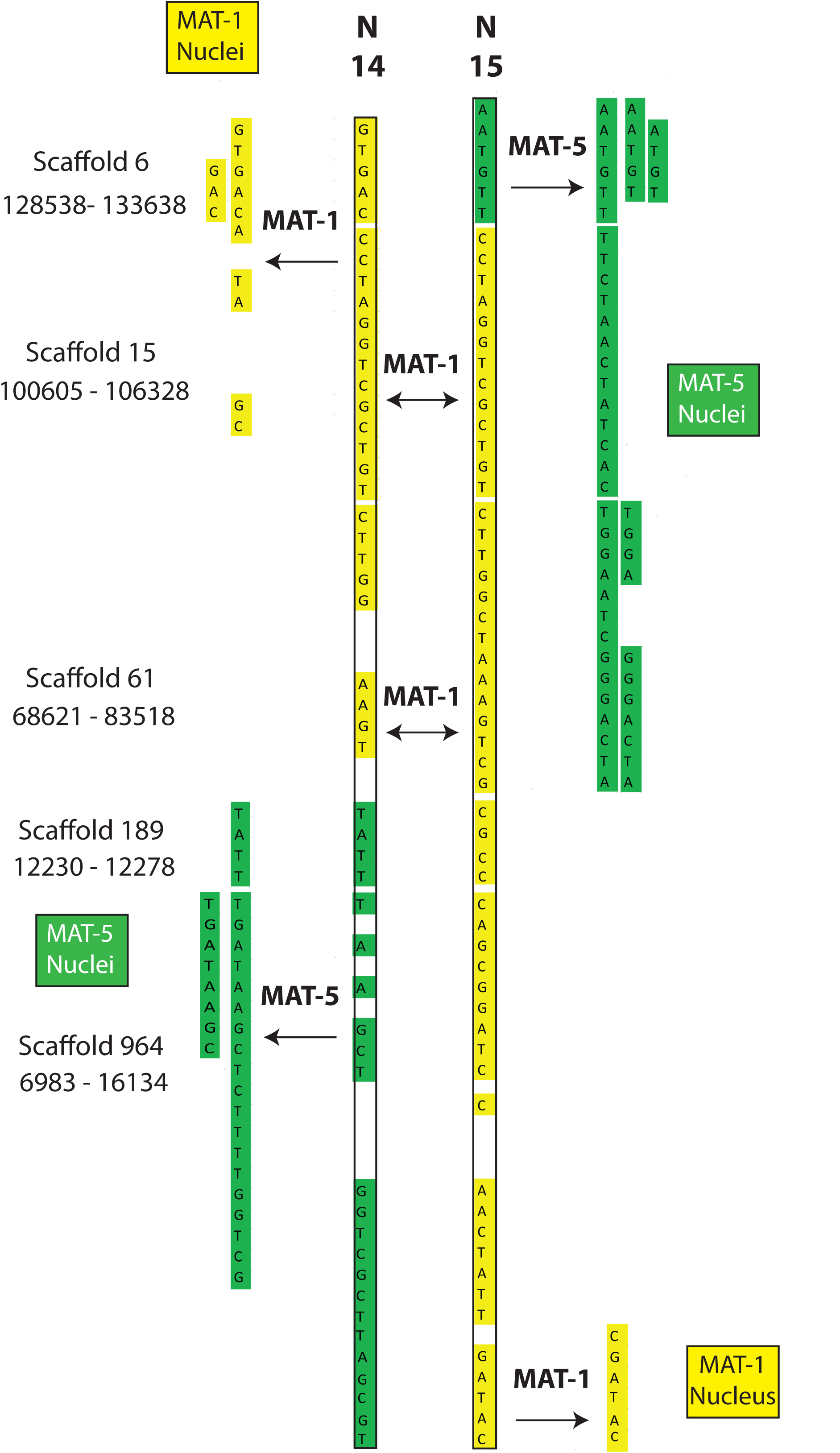
Examples of recombination involving two nuclei (SN14 and SN15) of the dikaryotic isolate SL1. The regions are found along homozygous regions present only once in the reference genome of SL1. The nuclei N14 and 15 from SL1 carry the MAT-1 locus (validated by PCR) and, accordingly, their sequenced genotypes are almost identical. In some cases, however, each nucleus swaps genotype with the opposite MAT locus (i.e. genotype becomes green).

Remarkably, by implementing the same read mapping method originally used by Chen et al. 2018, the exact genotypes originally found in ^1^ were recovered using the paired-end nucleus Illumina reads from A5. In all cases, the genotypes are linked with their respective MAT-locus and, importantly, all positions (16/16) with a read depth ranging from 1 to 4 produce the expected genotype (**Supplemental Table 3**).

We also aimed to determine if positions supported by two to four reads would result in a dramatic increase in recombination events. This would be expected if these SNP calls were mainly spurious. To do this, we sought evidence of inter-nucleus recombination along single copy, homozygous regions with minimum two reads supporting a SNP position. This analysis retrieved, respectively, 73, 168 and 31 recombinant blocks in SL1, A4 and A5, ranging from a minimum of 3 to a maximum of 21 contiguous SNPs (**Supplemental Table 4, Supplemental Table 5)**. Contrary to expectation, the number of recombinant blocks supported by less than five reads does remains within the same range – e.g. the number of blocks increases by 3% for A5 and 23% for A4). The larger block number increase seen in SL1 (36 to 73) simply reflects the low nucleus coverage of this isolate, which results from a genome reference that has more than twice the number of contigs compared to other dikaryotic isolates (despite being generated with longer mate-pair libraries and higher coverage) **(Supplemental Table 1)**.

Note that the putative recombination events found along regions supported by less than 5 reads (**Supplemental Table 4, Supplemental Table 5**) include cases where single nuclei swap genotypes back and forth along up to 60Kb (**Figures 2-3**); something that is difficult to explain based on low coverage alone. Lastly, many recombinant sites are also found in regions with high coverage ranging between 10 to 2321 (see a few examples in **Supplemental Table 6)**, thus evidence of inter-nucleus recombination is not necessarily linked with low coverage.

### Confirming that nucleus 07 (SL1) carries the MAT-5 locus would be compelling evidence against recombination in this nucleus

Using PCR and sequencing, Chen et al. 2018 found that Nucleus 07 of SL1 (SL1_07) carries the MAT-locus 1, even though its nuclear genotype mostly resembles those of co-existing MAT-5 nuclei. It was noted that Illumina reads covering the MAT-locus 1 locus are not present in the in SL1_07 single nucleus data, and that perhaps that specific sample was mixed-up.

Interestingly, we also note that: 1) Illumina reads that map the MAT-locus 5 are also absent from the SL1_07 data, even though their presence would provide compelling evidence against recombination in this nucleus and support of sample mix-up; 2) Other nuclei have no evidence of read mapping along the MAT-locus and all still had their MAT-locus identity properly confirmed by PCR/Sanger like SL1_07; 3) SL1_07 carries substantial evidence of recombination beyond the MAT-locus, particularly in the ALLPaths-LG assembly (see Supplementary file 7 in ^10^).

As such, the absence of sequencing reads covering the MAT-locus provides no evidence against the presence of recombination in the nucleus SL1_07.

### Conclusions

The aim of Chen *et al.* 2018 was not to make an inventory of inter-nucleus recombination events in AMF but rather to:

a. Validate the existence of a unique dikaryotic condition in some AMF isolates, whereby several thousands of nuclei that are copies of two parental genotypes co-exist in one large cell following plasmogamy between compatible homokaryons.
b. Identify the degree of nuclear diversity within the AMF mycelium, which is found to be always low in the genus *Rhizophagus.*
c. Detect evidence of inter-nucleus recombination in three dikaryotic isolates.

The recent comment paper on Chen *et al.* 2018 ^10,11^ focused on point (c). Yet, by re-analyzing the same single nucleus data with more stringent filters (single copy and homozygous sites with more than five reads supporting a SNP), dikaryotic isolates are still found to carry evidence of inter-nucleus genetic exchange. At a minimum, these findings confirm what is already known – i.e. co-existing nuclei in conventional dikaryotic cells (2 nuclei/cell) show footprints of recombination similar to those observed here ^8,9^. Within this context, to suggest that AMF does not undergo similar processes, one must assume that several thousands of nuclei from two parental genotypes can co-exist in the same cytoplasm for decades without undergoing genetic interactions.

Nonetheless, it is fitting to end on a cautionary note regarding the use of read mapping to genotype individual nuclei. Specifically, even though the present work validates previous findings and the methodology we used is appropriate to test inter-nuclear recombination, the work of Chen *et al.* 2018 also relies on sequence data that can vary dramatically in terms of coverage (as a result, for example, of multiple displacements during genome amplification, PCR bias during Illumina sequencing, or rare DNA cross-contamination ^12,13^). It also relies on a reference genome and pre-determined mapping thresholds that can all independently affect the analysis output.

Thus, like for any biological finding, it will be important for future studies to validate the presence of inter-nucleus recombination using alternative methods. To this end, plans are underway to sequence individual AMF nuclei using long-read sequencing technologies, and perform single nuclei genotyping using complete, phased genome references for all dikaryotic AMF strains. Lastly, producing a recombining progeny by crossing compatible strains will be key to demonstrate how/when sexual reproduction (meiosis) occurs in AMF.

## Methods

### Obtaining genotype files

For filtering and generating genotype files in **Supplemental Table 2 and Supplemental Table 5**, the original method described in Chen *et al.* 2018 was used with three modifications. First, the number of BLAST hits allowed is reduced to just one (from two) so that no duplicated region is taken into account. The second modification relates to the treatment of heterozygosity. Sites for individual nuclei that did not pass the 10-to-1 alternate to reference allele test based on Freebayes ^14^ SNP caller (hence forth referred to as “10-to-1”) are now removed, even in cases where their genotype is confirmed by homozygous nuclei, which was the approach originally used in ^10^. The final modification is extending the number of scaffolds surveyed to first 1000 scaffolds (from 100).

### Homokaryon low-coverage read analysis

To assess the fidelity of low coverage calls (Figure 1), homokaryon isolate A1, C2, *R. cerebriforme*, and *R. diaphanus*, are used. From the mapped BAM file of each single nucleus sequencing, we extracted positions from the first 10 scaffolds whose position have coverage of 1, 2, 3, 4, 5, 10, and 100. For each nucleus, positions with indel calls are filtered out. The 10-to-1 is also used on heterozygous positions. Finally, the percentage of homozygous mismatches are then calculated and collected across nuclei of each isolate before plotting in R via ggplot2, reshape2, grid, and grid_extra.

### Genotype identity

The goal of colour labels is to make it easier the observation of recombination footprints between nuclei. It is not to produce complete haplotypes. We assign genotype colour first based on parsimony using nuclei with PCR validated mating type: the mating type with more nuclei showing a particular genotype gets a colour assigned. If it is a tie, or there is no PCR-proven mating type, then colour that does not suggest new recombination is assigned (no change of colour down the column). If that fails, first genotype in that row to be *MAT*-A. The exception is SL1’s nucleus SN07 where in tied situations the colour corresponding to *MAT*-5 is assigned, which is consistent with its genotype clustering with other *MAT*-5 nuclei ^10^.

To score recombination events in scenario #1, a site is flagged as “recombining” if it starts to share one of two consecutive SNP with nuclei of the other MAT-locus along the same scaffold. For scenario #2, the same process is used, but a minimum of 3 consecutive SNP must be present.

For both scenarios, we count the number of events in each nucleus. For example, if 2 nuclei show recombination at the same location, the total number of events identified would be 2. Finally, in SL1’s SN07, we sometimes manually correct the colouring to highlight instances where it did not have recombination. This is purely for clarity only and does not affect the counting of recombination events.

### Obtaining read support of each position

To generate the read support for **Figure 1, Supplemental Table 1-6**, we opt to use bam-readcount (https://github.com/genome/bam-readcount; version 0.8.0). We used the original bam files from Chen *et al.* and queries for positions of interest. In the reanalysis of genotypes from Ropars *et al. 2016*, we used BLAST to identify the location of PCR products.

## Figure Captions

**Supplemental Table 1.** Genotype file with minimum coverage of 5, with heterozygous sites as defined in ^10,11^ removed, and single-copy regions. Yellow and Green colours highlight the two co-existing genotypes. Nuclei highlighted in yellow and green had their MAT locus validated by PCR. Nuclei with cells highlighted in light green carry a genotype that is mostly associated with green nuclei validated by PCR, while those with cells highlighted in orange carry a genotype that is mostly associated with yellow nuclei validated by PCR.

**Supplemental Table 2.** Recombination blocks (Scenario #2 obtained from genotype file with a at least five reads supporting a SNP in the first 1000 contigs. Based on **Supplemental Table 1**.

**Supplemental Table 3.** Confirmation of genotypes found by Ropars et al. 2016 Nature Microbiology using PCR and Sanger sequencing. F (A) == Reads supporting genotype (reads against genotype). Read depth is shown on the left columns for each position. Numbers in bold represent valid genotypes with very low read depth, i.e. 1 to 5.

**Supplemental Table 4.** Recombination blocks (Scenario #2 obtained from genotype file with a at least five reads supporting a SNP in the first 1000 contigs based on coverage < than 5. Based on **Supplemental Table 5.**

**Supplemental Table 5.** Genotype file with minimum coverage of 2, with heterozygous sites as defined in ^10,11^ removed, and based on regions found only once in the reference genome. Yellow and Green colours highlight the two co-existing genotypes. Nuclei highlighted in yellow and green in had their mating-type locus validated by PCR. Nuclei with cells highlighted in light green carry a genotype that is mostly associated with green nuclei validated by PCR, while those with cells highlighted in orange carry a genotype that is mostly associated with yellow nuclei validated by PCR.

**Supplemental Table 6**. Examples of recombination with high coverage. Nuclei ID are coloured for identification purposes. Yellow and dark green *MAT*-locus that is PCR verified. Yellow and Green colours highlight the two co-existing genotypes. Nuclei highlighted in yellow and green in had their mating-type locus validated by PCR. Nuclei with cells highlighted in light green carry a genotype that is mostly associated with green nuclei validated by PCR, while those with cells highlighted in orange carry a genotype that is mostly associated with yellow nuclei validated by PCR.

